# Improved Methodology for Studying Postnatal Osteogenesis via Intramembranous Ossification in a Murine Bone Marrow Injury Model

**DOI:** 10.1101/2024.10.24.620082

**Authors:** Marta Stetsiv, Matthew Wan, Shagun Prabhu, Rosa Guzzo, Archana Sanjay

**Author notes:** Equal contributions (Marta Stetsiv, Matthew Wan).

## Abstract

Long bone injuries heal through either endochondral or intramembranous bone formation pathways. Unlike the endochondral pathway that requires a cartilage template, the process of intramembranous ossification involves the direct conversion of skeletal stem and progenitor cells (SSPCs) into bone-forming osteoblasts. There are limited surgical methods to model this process in experimental mice. Here, we have improved upon a bone marrow injury model in mice to facilitate the study of bone repair via intramembranous ossification and to assess postnatal regulators of osteogenesis. This method is highly reproducible and user-friendly, and it allows temporal assessment of new bone formation in a short period (3-7 days post-injury) using µCT and frozen section histology. Furthermore, the contributions of SSPCs and mature osteoblasts can be readily assessed using a combination of fluorescent reporter mice and this intramembranous bone marrow injury model. In clinical contexts, intramembranous bone formation is relevant for healing critical size defects, stress fractures, cortical defects, trauma from tumor resections, and joint replacements.

**SUMMARY:** The murine bone marrow injury model is a useful tool to study postnatal osteogenesis through intramembranous ossification. This simplified surgical protocol describes the bone marrow ablation procedure for downstream assessment of new bone formation and cellular responses following injury.

## INTRODUCTION

Long bone injuries heal through the endochondral or intramembranous bone formation pathway. Unlike the endochondral pathway which requires a cartilage precursor step, bone repair via intramembranous ossification involves direct conversion of skeletal stem and progenitor cells (SSPCS) into bone-forming osteoblasts.^1^ Clinically, this process is relevant for bone healing in a variety of contexts, including critical size defects, stress fractures, cortical defects, distraction osteogenesis, trauma from tumor resections, and osseointegration of joint replacement implants.^2,3^ The mechanistic regulation of intramembranous bone regeneration remains relatively understudied compared to bone formation via the endochondral pathways. FDA-approved therapies to augment bone healing are hindered by limited clinical efficacy, high costs, and are associated with significant adverse effects.^4^

Mechanical bone marrow ablation provides a valuable model for studying intramembranous bone formation. This simple injury model allows for studying bone regeneration in the bone marrow without disrupting the cortical bone. Following BM ablation, the healing process involves distinct yet overlapping phases. The initial phase occurs in the first 1 to 5 days and involves clot formation and inflammation, where inflammatory cells and cytokines initiate healing. From days 3 to 14, the second phase is characterized by regeneration, including neovascularization, mesenchymal stem cell (MSC) migration and proliferation, osteoblastic differentiation, and woven bone formation. The final remodeling phase begins approximately 10 days after surgery, with bone tissue undergoing maturation and restructuring until the marrow is fully restored by day 56.^5^ This rapid healing timeline makes the BM ablation model ideal for studying early bone repair responses, particularly during the critical periods of inflammation, progenitor cell recruitment, and osteoblast differentiation. Using techniques like histology, flow cytometry, and quantitative µCT analysis, cellular responses and bone formation in early repair stages can be evaluated. Using the aforementioned techniques, the BM ablation model can provide insight into mechanisms of intramembranous bone regeneration and aid in identifying key therapeutic targets for enhancing bone healing.

### PROTOCOL

The following procedures were performed with approval from the University of Connecticut Health Center Institutional Animal Care and Use Committee (IACUC). All surgeries were performed under sterile conditions as outlined by the NIH guidelines. Pain and risk of infections were managed with proper analgesics and antibiotics to ensure a successful outcome.

#### 1. PREPARATION OF SURGICAL AREA AND INSTRUMENTS

1. Prepare the surgical procedure room by disinfecting working surfaces with 10% Lysol or other clinical grade disinfectant cleaner. NOTE: The surgeon should wear appropriate personal protective equipment (PPE), including sterile disposable gloves, gown, and mask.
2. Prepare a heating pad covered with a sterile surgical pad. Gather and arrange all sterile instruments and reagents **(Table of Materials)** with convenient access.

#### 2. PRE-OPERATIVE ANESTHESIA

1. Weigh all mice before surgery. Calculate the volumes of analgesic to be used based on the available formulation of buprenorphine.
2. Assemble an animal anesthesia unit within the sterile working space, which includes an induction chamber and a nose-cone assembly. Set oxygen output to 2 L min^-1^ and isoflurane to 1.5-3% (v/v) depending on mouse age and weight.
3. Once the mouse is sedated, remove the animal from the induction chamber and place it on a sterile surgical pad in the supine position. Place the nose cone over the head. Check that the mouse is adequately anesthetized by gently pinching the toe. Ensure there is no hindlimb reflex response before proceeding.
4. Prior to surgery, inject ½ of the total dose of buprenorphine analgesic via subcutaneous route. For regular buprenorphine, the total dose is 0.05-0.1 mg/kg, for extended-release formulation, the total dose is 3.25 mg/kg.

#### 3. SURGICAL PROCEDURE DESCRIPTION

This surgical procedure is optimized for C57BL/6 male and female mice aged 10 weeks to 24 weeks.

NOTE: The procedure can be performed on younger or older mice of either sex

1. Fill a 3 mL syringe equipped with a 26G needle with sterile saline. Place this to the side.
2. Remove the fur from the hindlimb on which procedure will be performed with electric or battery-operated clippers (#40 blade). Using sterile cotton tipped applicators, apply an iodine-based scrub (10% Povidone iodine or similar antiseptic) to the surgical site. Scrub the surgical site starting at the center of the knee and making a circular sweep outward. Allow the site to dry.
3. Place the mouse on its back and flex the knee of the operative leg. Hold the leg of the mouse in the non-dominant hand such that the knee is bent over the middle finger, the second finger rests on the femur, and the thumb rests on the tibia.
4. Make a small, shallow horizontal incision just below the patella using a scalpel. NOTE: This step improves visualization of the insertion site. With experience, this step can be skipped.
5. Locate the joint space by placing the needle horizontally across the bent knee and feeling for the space between the femur and tibia. Using this anatomical site to guide you, take a 25G needle and manually drill into the distal femur from the knee toward the proximal side. Needle Insertion may be through or around the patella.
6. X-ray the mouse to ensure the needle has been placed properly in the medullary cavity, as close to center as possible. [**Figure 1**]
7. Remove the 25G needle and immediately insert a 26G needle into the same hole within the medullary canal. Ream the sides of the medullary cavity until it feels smooth.
8. Remove the 26G needle and replace the 26G needle connected to the saline syringe. Flush the cavity until the emerging fluid runs clear. This typically requires 1.5 to 2 mL of saline. Remove the syringe and needle and safely dispose into a sharp’s container. NOTE: There should not be resistance when pushing the fluid. If so, re-insert the needle to ensure the needle is within the cavity. It may be necessary to pull the needle out halfway.
9. Close the skin wound using a topical adhesive suture.

**Figure 1.**
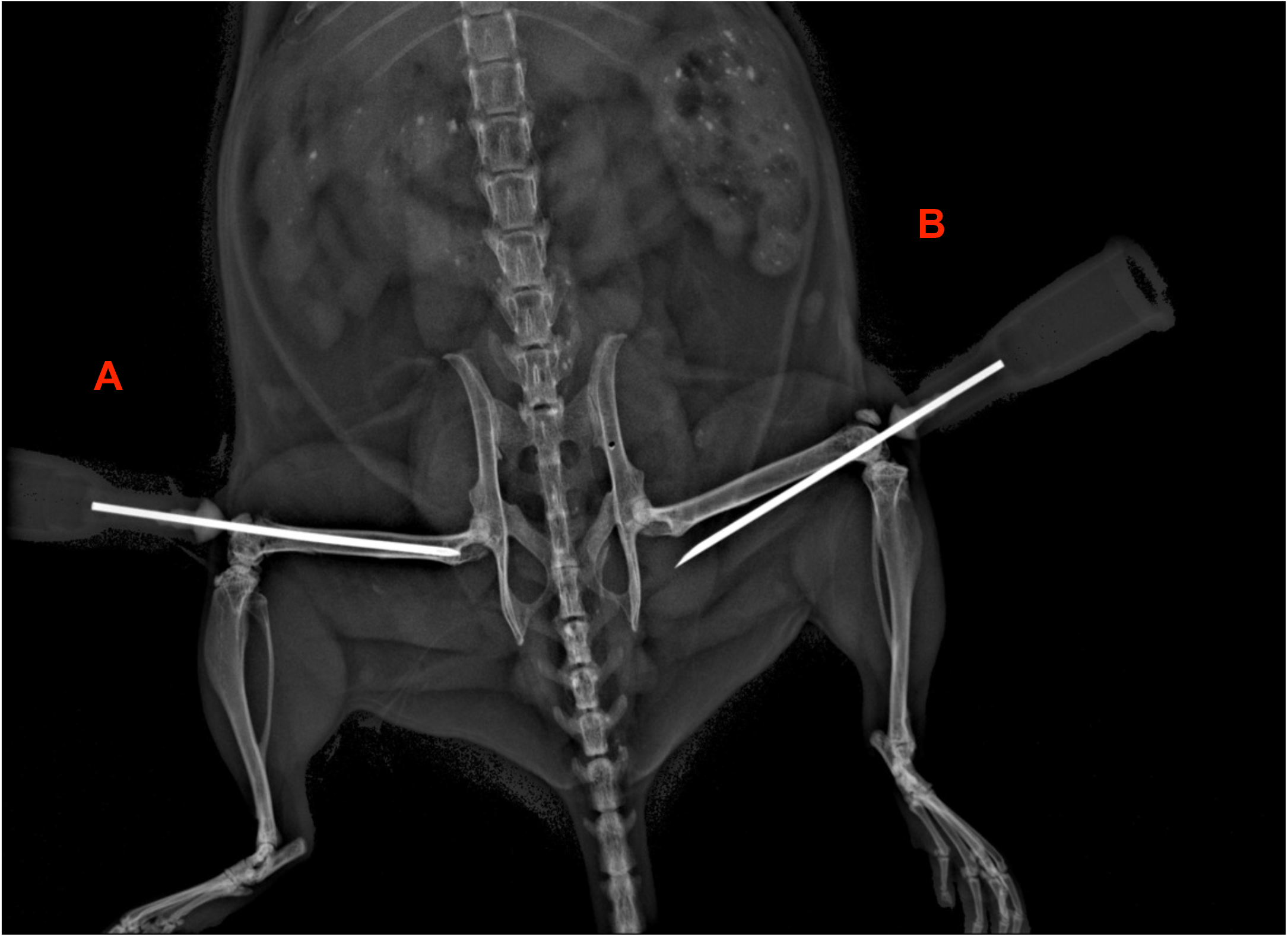
Examples of needle placement for BM ablation procedure. X-ray depicting bilateral BM ablation procedure, where the needle is successfully placed in the center of the femur bone **(A)**. Example of an unsuccessful needle placement, where the needle has pierced through the bone **(B)**.

#### 4. POSTOPERATIVE CARE

1. Immediately following surgery, administer the remaining half dose of extended-release buprenorphine using dosing from section 2.3. NOTE: Extended-release Buprenorphine provides pain relief for up to 72 hours.
2. Monitor the animals on the heating pad for signs of normal, unobstructed breathing until they awaken from surgery. Once ambulatory, return mice to their cage.
3. Continue to monitor the mice once per day for 3 days after the surgery to ensure they are healing properly and have full mobility, watching for signs of pain.

#### 5. PROCESSING BONES FOR μCT SCANNING AND ANALYSIS

1. Euthanize the mice according to IACUC institutional policy.
2. Harvest the femurs by peeling away the skin around the hindlimbs and remove the entire limb at the hip joint.
3. Sandwich each limb between two sponges in an embedding cassette. Transfer the bones in their cassettes to a 10% formalin solution and fix at 4 °C for 5-7 days considering the age and size of the mouse. NOTE: Both over fixation and under fixation result in poor marrow resolution.
4. After fixation, wash the cassettes in PBS for 5 minutes 3 times.
5. Remove the limbs from the cassettes and trim away muscle using a scalpel as needed. NOTE: Depending on downstream applications, rough handling can disrupt the periosteal layer of bone.
6. Separate the femur at the knee by carefully cutting through the ligaments using small surgical scissors or a scalpel. The tibias can be used for other applications or discarded.
7. Transfer the femurs to 1.7 mL or 15 mL conical tubes into a 70% ethanol solution. Bones can remain in 70% ethanol at 4 °C until ready for µCT scanning.
8. The femurs can be scanned and assessed for trabecular parameters, producing qualitative and quantitative data.

#### 5. PROCESSING BONES FOR FROZEN SECTIONING AND HISTOLOGY

NOTE: Bones can be processed for frozen sectioning and histology after undergoing µCT scanning. The scanning does not affect fluorescent labeling or subsequent antibody staining.

1. Transfer the bones from 70% ethanol to PBS and wash 3 times, 5 minutes per wash.
2. Incubate in a series of sucrose solutions from 10%, 20%, to 30% in PBS at 4°C overnight at each concentration. Bones may be stored in 30% sucrose for up to 5 days at 4 °C or in this solution at -20 °C for 6-12 months until ready for embedding. NOTE: Using a sucrose gradient results in better tissue penetration. After step 1, bones may be placed directly into 30% sucrose solution overnight. Embed bones in OCT using cryoembedding molds. Store in –80 °C.
3. Section embedded bones using the cryofilm tape method with cryostat temperature set between -24 °C and -26 °C.
4. Sections on cryofilm can be adhered to a superfrost slide using optical adhesive and cured until hardened using a UV crosslinker.

#### 6. PROCESSING FOR FLOW CYTOMETRY

NOTE: This protocol is compatible with a variety of antibodies. For characterizing skeletal stem and progenitor cells, pooling 3-4 bones together is recommended as these are rare cell populations in the bone marrow.

1. Prepare FACS Staining Media (FSM), to make 1000 mL solution of FSM:
  a. 100 mL HBSS
  b. 10 mL HEPES
  c. 20 mL FBS
  d. 870 mL dd H2O.
2. Before harvesting tissue, fill petri dishes with 4-5 mL of sterile PBS without calcium and magnesium.
3. Prepare the digestion media
  a. For every 4-6 femurs prepare digestion mixture in 10 mL PBS containing
  b. 0.005 g collagenase P
  c. 0.02 g hyaluronidase
  d. Mix gentle vortexing
  e. pass the mixture through a 0.22 µm filter
4. Harvest bones on post-surgery Day 3 as described in section 4.1. Remove all muscle- and connective tissue with a scalpel blade.
5. Fill a petri dish with 5 mL of digestion media. Place 4-6 bones in the digestion media. Chop each bone finely, making the first cut longitudinally and laterally. Transfer tissue to a 14 mL round-bottom tube and seal with parafilm. Cover tubes in aluminum foil.
6. Shake at 37 °C for 20 minutes by keeping the tube horizontally on the shaker
7. Remove from shaker. Cells will be clumped at the bottom. Gently collect the digest without disturbing the cell at the bottom, and pass the digest through a 70 µm filter. Collect the filtrate in a new 50 mL tube containing 10 mL of cold FSM and keep it on ice
8. To the cells add the remining 5 ml of the digestion mix. Seal again with parafilm and cover with aluminum foil. Shake the tube as described above. Repeat steps 6.6-6.7.
9. After the 20-minute incubation at 37 °C, transfer the digest to the same tube containing the digest from step 7, passing through a 70 µm filter with another 10 mL of FSM.
10. Centrifuge the suspension at 274 x g at 4 °C. Aspirate the supernatant and collect the cell pellet. Dislodge the pellet by gently tapping the tube. NOTE: *g* force based on a rotor radius of 170 mm. Please check your rotor radius and adjust speed accordingly.
11. Perform the red blood cells lysis.
12. Add 1 mL of ACK buffer. Keep all reagents and cells on ice.
  a. .Add 1 mL of ammonium-chloride-potassium (ACK) lysing buffer.
  b. Incubate the cells in ACK for about 5 minutes at room temperature. Do not exceed 5 minutes to avoid overly lysing the cells.
  c. Add 9 mL of FSM. Centrifuge at 274x *g* for 5 minutes at 4 °C.
  d. Aspirate the supernatant. Break up the cell pellet by gently raking the tube on a tube rack.
  e. Add 10 mL of FSM (volume for 4 bones, can be adjusted based on how many bones are combined) and resuspend
  f. Take an aliquot of each sample and add propidium iodide (PI). Count live and total cells with a cell counter.
13. Proceed with antibody staining of samples using desired markers for flow cytometry analysis.^6-8^

### REPRESENTATIVE RESULTS

The success of the BM ablation procedure was validated by X-ray imaging while the 26G needle was inserted into the medullary cavity (**Figure 2A**). When X-ray imaging indicated that the needle was inserted at an improper angle or penetrated through the bone, the needle was re-inserted at the proper angle and re-imaged to confirm accurate placement. When a successful position could not be attained, the animal was excluded from analysis. While radiographic 2D imaging can confirm that the needle was inserted within the BM cavity, 3D imaging is necessary to confirm the precise needle path. The injury extent and location were further assessed upon tissue harvest by qualitative µCT analysis of the femurs (**Figure 2B**). At 7 days post injury there was a rapid and robust regeneration of trabecular bone, as evidenced by a seven-fold increase in bone volume fraction and doubling in trabecular number (**Figure 2C-E)**, as well as increase in trabecular thickness and a decrease in trabecular spacing **(Figure 2F-G)**.

**Figure 2.**
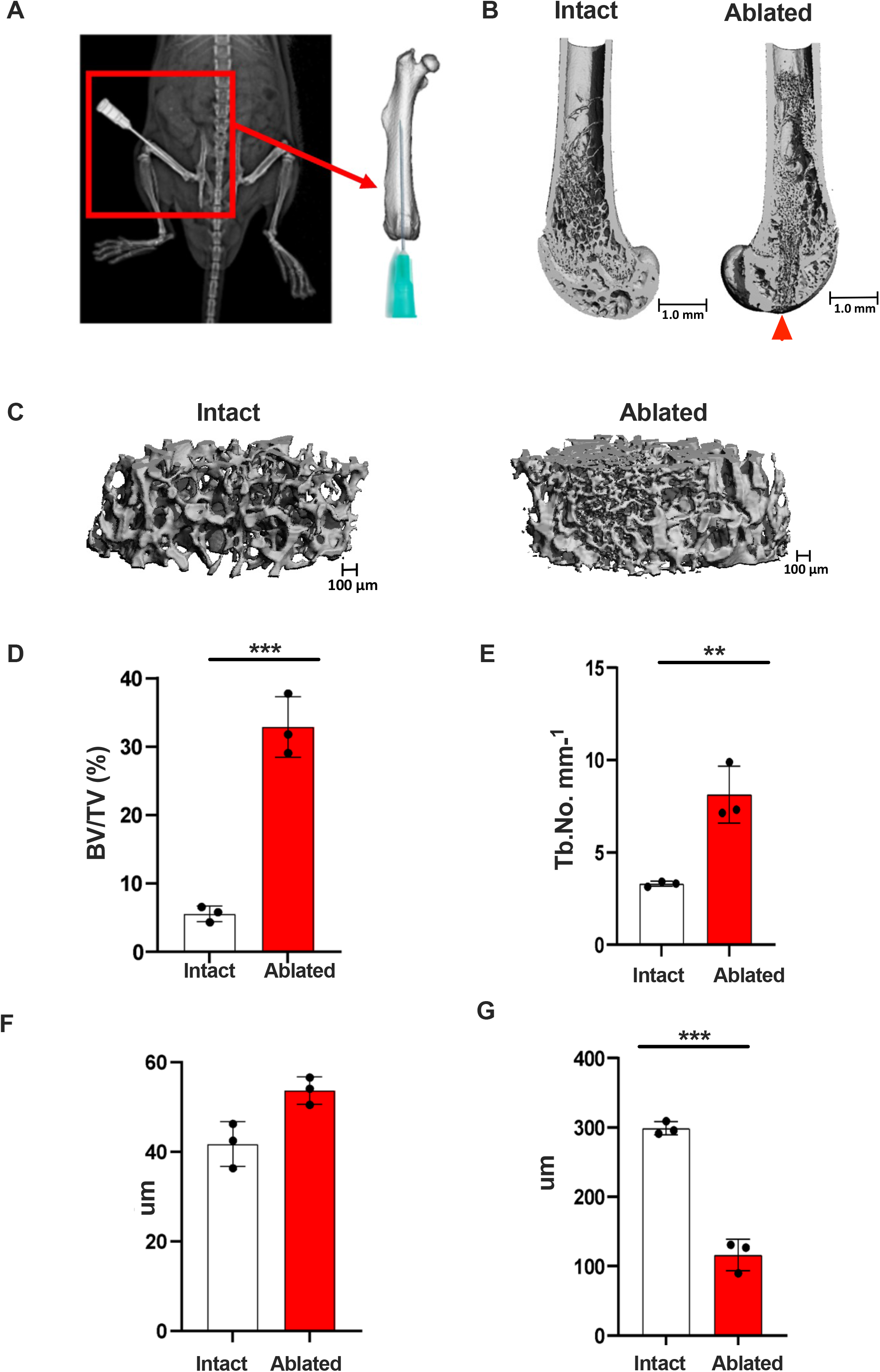
Qualitative and quantitative µCT analysis of intramembranous bone formation at 7 days post mechanical bone marrow ablation. **A)** 26G needle placement (right femur) is confirmed using X-ray imaging. **(B)** µCT images of intact and ablated femurs. The ablated femur shows the needle path through the center of the bone from the proximal end (arrow shows insertion point). **(C)** Micro-CT images of a section of trabecular bone cores from intact and ablated femurs. **(D-G)** Quantification of bone volume/total volume (BV/TV), trabecular number, trabecular thickness, and trabecular spacing in intact vs ablated femurs. n = 3; mean ± SD; * p < 0.05

General architecture of the growth plate and bone marrow cavity is shown by Toluidine Blue staining of intact and ablated bone **(Figure 3A)**. An increase in bone formation after surgery was further confirmed qualitatively via von Kossa staining (**Figure 3B**). To confirm that this injury heals via intramembranous ossification, Safranin O/Fast Green staining was performed on frozen sections of intact and ablated femurs to visualize cartilage. This staining shows cartilage in the intact growth plate, but no cartilage formation in response to injury (**Figure 3C**).

**Figure 3.**
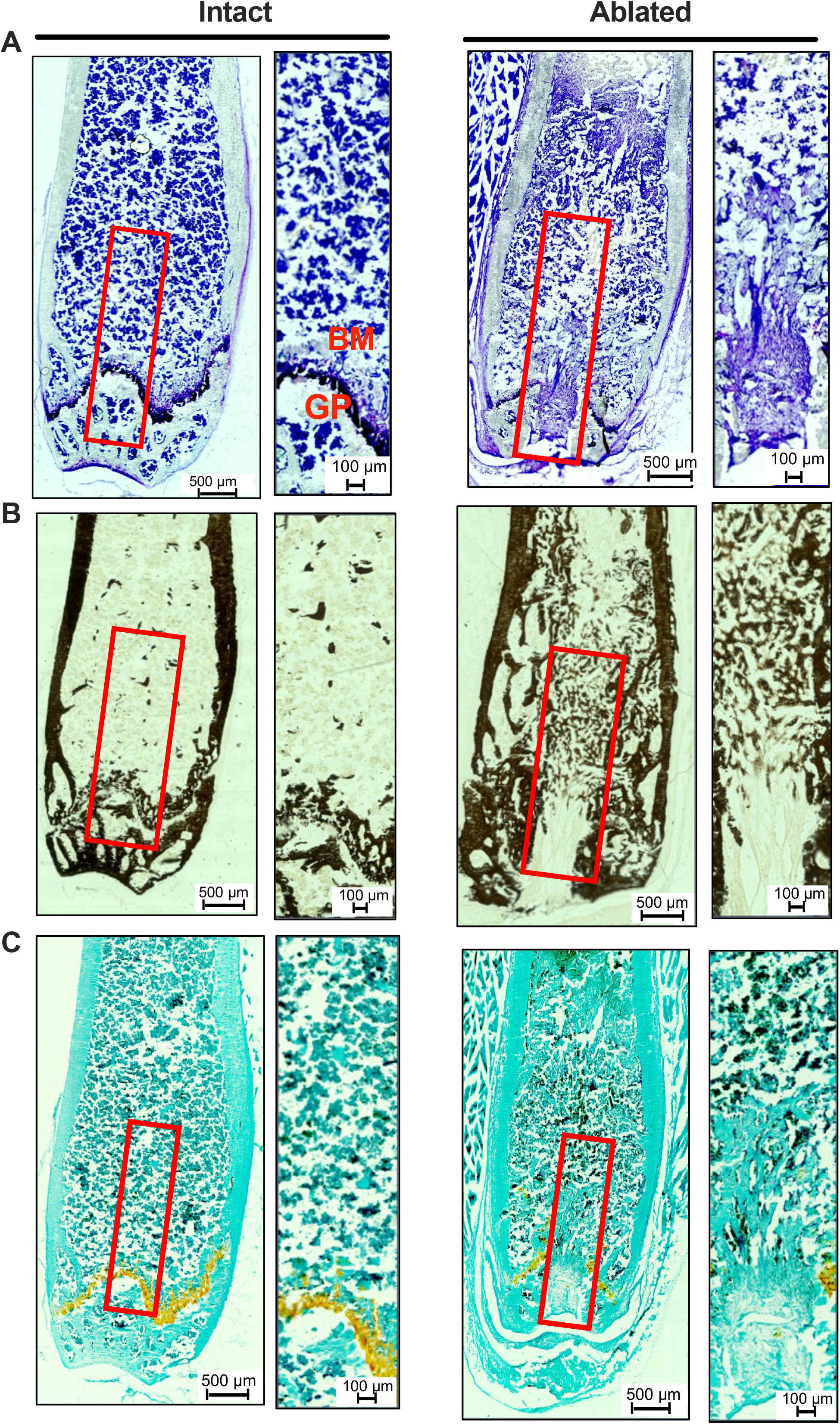
Qualitative assessment of intramembranous bone formation by histology. **(A)** Frozen sections (9 µm) of intact and ablated femurs stained with Toluidine Blue to visualize general architecture. Outlined areas in the path of the needle (red) shown at higher magnification to the right. **(B)** von Kossa mineral staining showing robust bone formation in the ablated femur compared to intact control. Outlined areas (red) shown at higher magnification to the right. **(C)** Safranin O/Fast Green staining showing cartilage in the growth plate (orange) in the intact bone (left) but not present in the ablated femur (right). Outlined areas (red) shown at higher magnification. GP = growth plate; BM = bone marrow.

Another approach to assess *de novo* bone formation in this injury model is calcein labeling. Mice received IP calcein injections at 20 mg/kg 24 hours before a Day 7 harvest (**Figure 4A**). Calcein mineral labeling was visualized using fluorescent microscopy, which shows an increase in calcein label in the ablated femur following the path of the needle injury. (**Figure 4B**).

**Figure 4.**
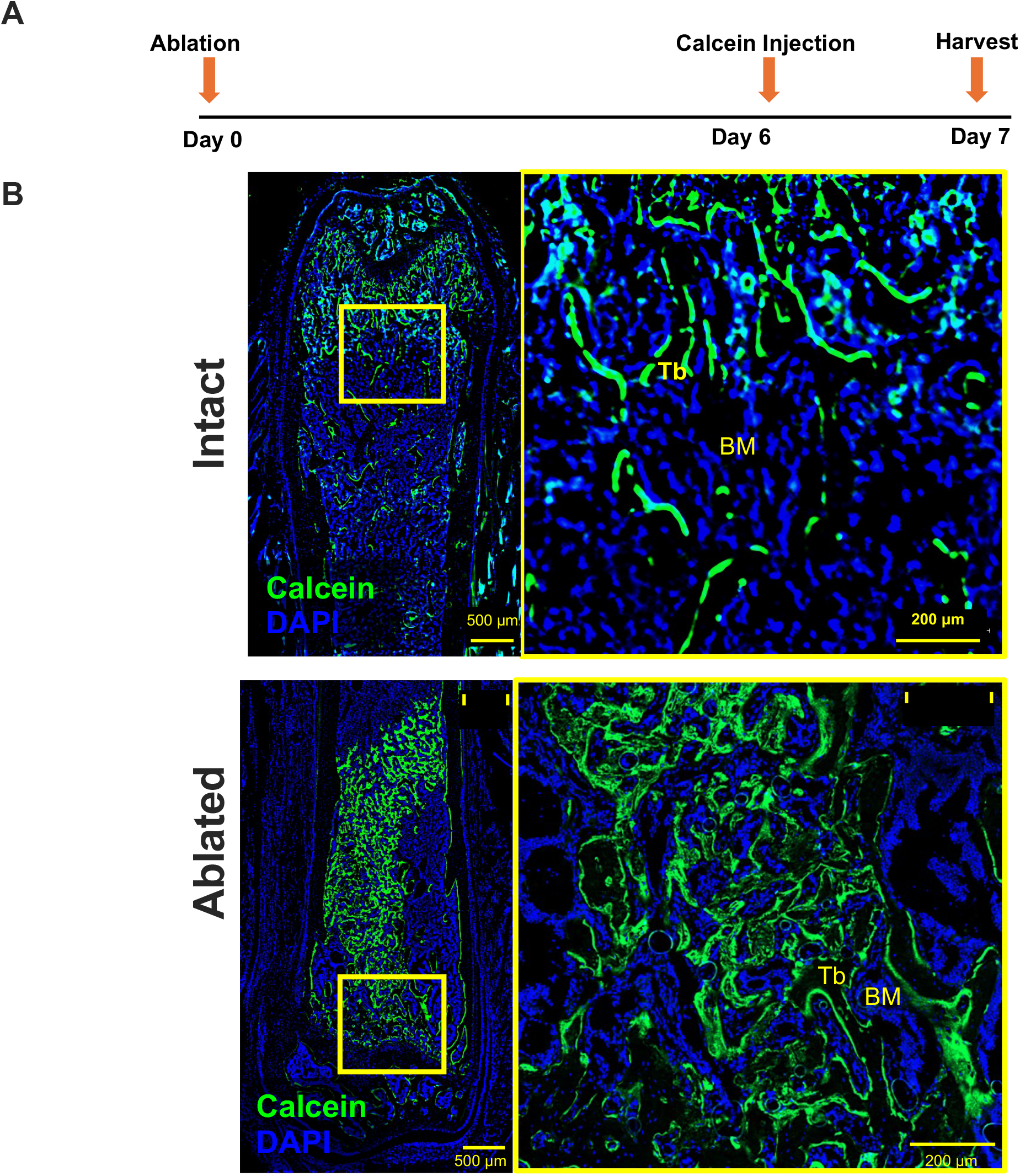
Tracking new bone formation using calcein labeling. **(A)** Schematic of experimental design. **(B)** Fluorescent calcein labeling (green) and nuclear DAPI label (blue) are shown in intact (left) and ablated (right) femurs. High magnification images depicting the outlined areas (red box) show new bone formation along the path of the needle in the ablated bone. BM = bone marrow; Tb = trabeculae.

This model can also be used with Cre reporter mice to study skeletal progenitor responses in intramembranous bone regeneration. We performed unilateral BM ablation in 24-week-old Prrx1-Cre; Ai9^tdTomato^ mice. Paired related homeobox 1 (Prrx1) marks early limb bud progenitor cells. The femurs were harvested 7 days after surgery and showed an increase in tdTomato+ cells compared to the intact control (**Figure 5A-B**). Other Cre mouse lines may be used with the BM ablation model to study bone marrow specific skeletal stem and progenitor cell responses to injury.

**Figure 5.**
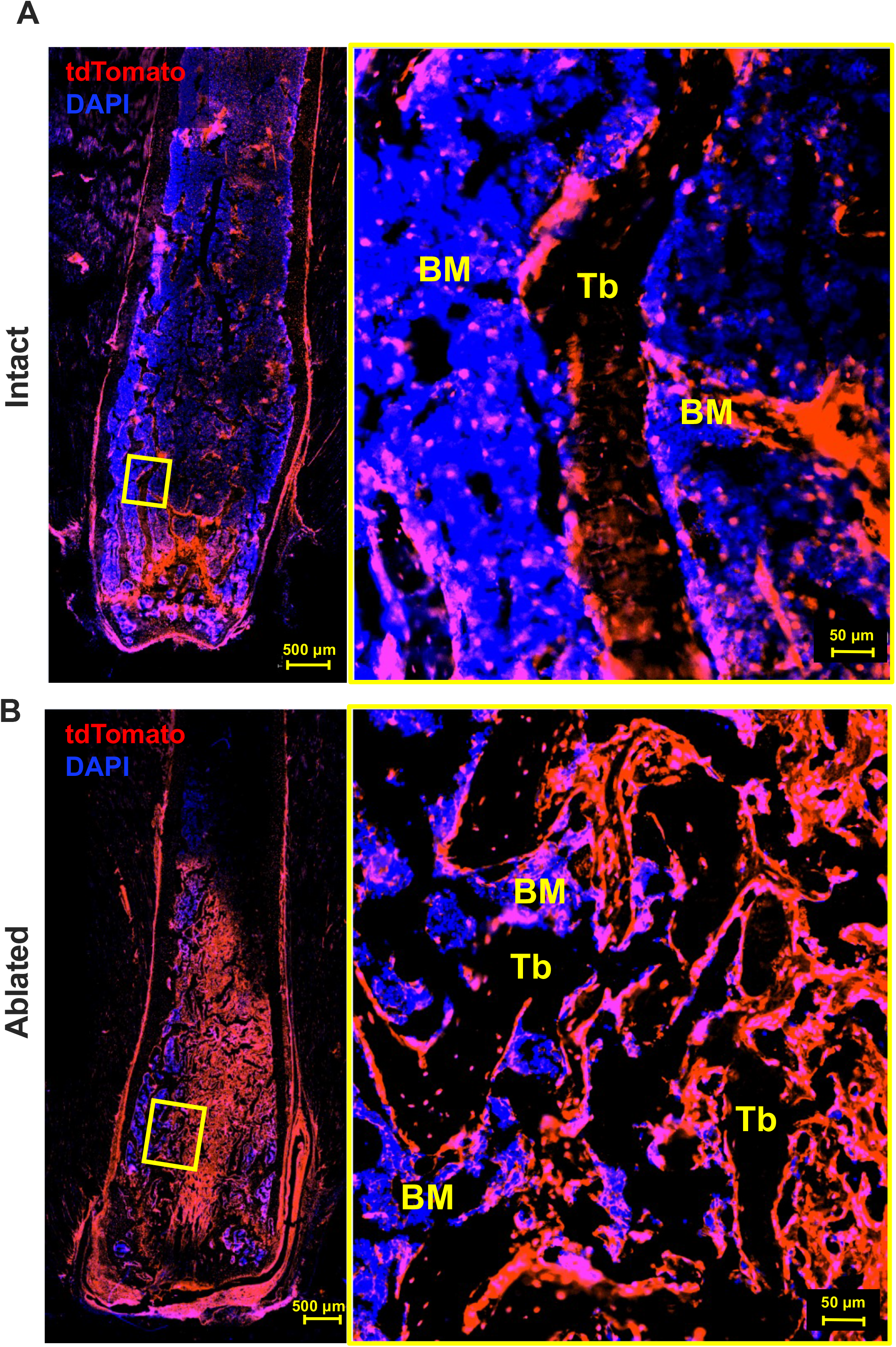
Fluorescent imaging showing increased numbers of tdTomato+ cells in response to BM ablation using Prrx1-Cre reporter mice. **(A)** Localization of tdTomato+ cells in an intact femur. Nuclei were counterstained with DAPI. Outlined area (yellow) shows tdTomato+ cells at high magnification. **(B)** A representative image showing a robust increase in tdTomato expressing cells is seen in the needle’s path, with an enrichment in cells lining the trabeculae, in a femur harvested 7 days after BM ablation. BM = bone marrow; Tb = trabeculae.

Together, our data demonstrate the robustness of the BM ablation model and the many tools that can be utilized with it to assess de novo bone formation and marrow specific cellular responses to injury.

### TABLE OF MATERIALS

Supplemental Table 1

## DISCUSSION

The goal was to establish a reliable and reproducible method to study the molecular and cellular processes governing bone repair, focusing on the direct conversion of skeletal stem and progenitor cells (SSPCs) into bone depositing osteoblasts in the bone marrow space/cavity. The mechanical bone marrow ablation method described here is an optimized and validated protocol for bone marrow ablation by mechanical injury, which allows for the investigation of intramembranous bone formation. Several steps in this protocol are critical to ensure robust bone formation post injury and reproducibility of the technique. First, is the precise insertion of the needle into the femoral medullary cavity. Misalignment in this step can lead to incomplete marrow ablation and damage to surrounding tissues, leading to outcome variability. To ensure proper needle placement, X-ray imaging must be utilized to visualize needle placement before reaming the marrow cavity. Another critical step is flushing the bone marrow cavity with saline to remove BM remnants and unpack the marrow creating space for subsequent new bone formation. Modifications to this protocol may be needed depending on the specific research question, mouse strain, genotype, or mouse age and weight.^9,10^ The size of the needles and volume of saline flush may be adjusted based on the animal size and age. Common issues related to this surgical technique may include resistance when flushing the marrow cavity or animals not waking up post-surgery. If resistance to saline flushing is encountered, the needle can be repositioned or pulled out halfway. If this does not resolve resistance to flushing, marrow reaming may need to be repeated to ensure the BM cavity is clear of bone remnants.

Intramembranous bone formation has been studied using various injury models.^11-13^ Among commonly used models are the calvarial defect model and the critical size defect model.^12,14^ The calvarial defect model allows for ease of access and the ability to test a variety of biomaterials to assess osteogenic potential.^15^ However, this model includes flat bone healing, which differs from long bones, where the mechanical environment and cellular dynamics are distinct. The critical size defect model involves creating a defect in the femur or tibia that is too large to heal spontaneously, mimicking clinical scenarios like trauma or tumor resection, which require intervention to achieve healing. Significant limitations of this model are the invasive surgical procedure to create critical size defect, requirement of specialized screws and plates, and extended healing period, which can be a disadvantage to studying early cellular events in bone repair.^13,16^ Additionally, endochondral ossification can sometimes occur in healing critical size defects.^17^ In contrast to these injury models, the mechanical BM ablation model provides a more controlled environment and reproducible injury responses that facilitate the study of intramembranous bone formation, specifically in long bones. This model targets the intramembranous pathway (as evidenced by a lack of cartilage formation visualized by Safranin O staining in our examples), allowing for more precise studies of SSPC contributions to bone healing. This is particularly beneficial for studies aimed at understanding the early stages of bone regeneration, where the dynamics of SSPC recruitment and differentiation are most critical. Additionally, using fluorescent reporter mice with this model enables a detailed analysis of SSPC contributions.

Another advantage of this model is the range of downstream applications and imaging modalities that can be used to assess qualitative and quantitative bone formation and cellular responses to injury. Early cellular responses can be assessed on Day 3 after the surgery, using EdU labeling for proliferation, flow cytometry for skeletal and immune cell surface markers, RNAscope for spatial visualization of gene expression patterns, and single cell RNA sequencing to evaluate responses of different cell types. New bone formation can be assessed as early as Day 7 after the procedure, using dynamic histomorphometry, qualitative and quantitative µCT analysis, frozen section histology, and calcein labeling. Additionally, this model can be used to study bone resorption starting on Day 10 post-surgery and TRAP staining to visualize osteoclasts.

While this injury model is highly effective for studying intramembranous bone formation, there are some limitations. One such limitation is the dependence on surgical skill of the surgeon performing the technique, which can introduce variability in outcome. However, the learning curve for this protocol is relatively small and is aided by validation of needle placement using X-ray imaging. Another key challenge is the inherent heterogeneity of the bone marrow microenvironment. The BM comprises multiple distinct niches, including hematopoietic, stromal, vascular, and osteoblastic compartments, which may participate in crosstalk to regulate the homeostasis of the bone and marrow.^18,19^ This can make it difficult to distinguish specific contributions of individual compartments and cell types to the bone repair process. Additionally, the surgical procedure in this model disrupts the growth plate, which harbors chondrocyte-derived progenitors that contribute to bone development. A small subset of these chondrocytes can transform into osteoblasts and marrow stromal cells, even at the postnatal stages.^20-22^ This could complicate interpretation of bone marrow derived progenitor responses to injury. Particularly, disruption of the growth plate can be an issue in younger mice, where there is higher activity of progenitor cells in a growing bone.

The BM ablation injury model has significant implications for research areas focused on bone repair and regeneration. It is particularly useful for studies investigating the contributions of SSPCs in bone healing and novel therapeutic strategies to target these cells. This model can be used in translational studies to evaluate the efficacy of novel small molecules and drugs in promoting bone healing in a controlled setting. Clinically, insights gained from the use of this model can contribute to the development of improved treatments for conditions requiring intramembranous bone formation, such as fracture non-union, large bone defects, bone loss associated with tumor resection, or osseointegration in joint replacement.

In summary, BM ablation injury model is a valuable and versatile tool for studying intramembranous bone formation. While there is a small learning curve in the surgical technique, with experience the method provides reproducible outcomes. Insights into mechanisms of bone repair using this model make it a powerful tool for orthopedic research.

## Supporting information

Supplemental Table 1: Materials List

## ACKNOWLEDGMENTS

Funding was provided by NIH/NIDCR R01-DE030716-01 and NIH/NIAMS R01-DE030716-01S1 to AS, NIH/NIAMS R01AR080131 to RG, and UConn REP Convergence Grant to RG and AS.

## REFERENCES

1. Bahney, C. S. et al. Cellular biology of fracture healing. Journal Orthopaedic Research 37, 35–50 (2019).

2. Ko, F. C. & Sumner, D. R. How faithfully does intramembranous bone regeneration recapitulate embryonic skeletal development? Developmental Dynamics 250, 377–392 (2021).

3. Patt, H. M. & Maloney, M. A. Bone marrow regeneration after local injury: a review. Exp Hematol 3, 135–148 (1975).

4. Katagiri, T. et al. Bone morphogenetic protein-induced heterotopic bone formation: What have we learned from the history of a half century? Japanese Dental Science Review 51, 42–50 (2015).

5. Wise, J. K. et al. Temporal Gene Expression Profiling during Rat Femoral Marrow Ablation-Induced Intramembranous Bone Regeneration. PLoS ONE 5, e12987 (2010).

6. Jeffery, E. C., Mann, T. L. A., Pool, J. A., Zhao, Z. & Morrison, S. J. Bone marrow and periosteal skeletal stem/progenitor cells contribute distinct to bone maintenance and repair. Cell Stem Cell 29, 1547-1561.e6 (2022).

7. Loopmans, S., Stockmans, I., Carmeliet, G. & Stegen, S. Isolation and in vitro characterization of murine young-adult long bone skeletal progenitors. Front. Endocrinol. 13, 930358 (2022).

8. Chan, C. K. F. et al. Identification and Specification of the Mouse Skeletal Stem Cell. Cell 160, 285–298 (2015).

9. Moran, M. M. et al. Intramembranous bone regeneration differs among common inbred mouse strains following marrow ablation. Journal Orthopaedic Research 33, 1374–1381 (2015).

10. Moran, M. M. et al. Intramembranous bone regeneration in diversity outbred mice is heritable. Bone 164, 116524 (2022).

11. Lybrand, K., Bragdon, B. & Gerstenfeld, L. Mouse Models of Bone Healing: Fracture, Marrow Ablation, and Distraction Osteogenesis. CP Mouse Biology 5, 35–49 (2015). 12.

12. Cooper, G. M. et al. Testing the Critical Size in Calvarial Bone Defects: Revisiting the Concept of a Critical-Size Defect: Plastic and Reconstructive Surgery 125, 1685–1692 (2010).

13. Yu, Y., Bahney, C., Hu, D., Marcucio, R. S. & Miclau Iii, T. Creating Rigidly Stabilized Fractures for Assessing Intramembranous Ossification, Distraction Osteogenesis, or Healing of Critical Sized Defects. JoVE 3552 (2012).

14. Wu, X. et al. Enhancing calvarial defects repair with PDGF-BB mimetic peptide hydrogels. Journal of Controlled Release 370, 277–286 (2024).

15. Stanton, E. et al. A Murine Calvarial Defect Model for the Investigation of the Osteogenic Potential of Newborn Umbilical Cord Mesenchymal Stem Cells in Bone Regeneration. Plastic & Reconstructive Surgery (2023).

16. Manassero, M. et al. Establishment of a Segmental Femoral Critical-size Defect Model in Mice Stabilized by Plate Osteosynthesis. JoVE 52940 (2016).

17. Petersen, A. et al. A biomaterial with a channel-like pore architecture induces endochondral healing of bone defects. Nat Commun 9, 4430 (2018).

18. Xiao, X. et al. Spatial transcriptomic interrogation of the murine bone marrow signaling landscape. Bone Res 11, 59 (2023).

19. Yu, V. W. C. & Scadden, D. T. Heterogeneity of the bone marrow niche. Current Opinion in Hematology 23, 331–338 (2016).

20. Matsushita, Y., Ono, W. & Ono, N. Growth plate skeletal stem cells and their transition from cartilage to bone. Bone 136, 115359 (2020).

21. Mizuhashi, K. et al. Resting zone of the growth plate houses a unique class of skeletal stem cells. Nature 563, 254–258 (2018).

22. Muruganandan, S. et al. A FoxA2+ long-term stem cell population is necessary for growth plate cartilage regeneration after injury. Nat Commun 13, 2515 (2022).

