## Supplemental Table 1: Materials List for "Improved Methodology for Studying Postnatal Osteogenesis via Intramembranous Ossification in a Murine Bone Marrow Injury Model"

| Name of Material/ Equipment | Company | Catalog Number | Comments/Description |
|---|---|---|---|
| 10% Formalin Solution | Sigma-Aldrich | HT501128 | |
| 25G 0.625 in Air-tite EZ Flo Hypodermic Needle | Ait Tite/ Fisher Scientific | 14-817-239; ref #NEZ25058 | |
| 26G 1/2" Disposable Hypodermic Needles | Exel International/ Fisher Scientific | 14-840-83; ref #264 | Any 26G needles may be substituted, must be detachable from syringe |
| 3 mL Sterile Syringe | Fisherbrand/ Fisher Scientific | 14-955-457 | |
| AB Sterile Pad | McKesson/ Fisher Scientific | NC0593755 | Alternatively can use autoclavable absorbant pads |
| Acetic Acid, Glacial | Millipore Sigma | AX0073 | Needs to be diluted to 1% |
| Ammonium-Chloride-Potassium (ACK) Lysing Buffer | Gibco/ Thermo Fisher Scientific | A1049201 | Used for red blood cell lysis in preparation for flow cytometry |
| Anesthesia Induction Chamber 2L | VetEquip | 941444 | |
| Aseptic Control Envirocide Disinfectant Cleaner | Aseptic Control Products/ Fisher Scientifi | 500432411 | Can use any clincial grade disinfectant solution |
| Betadine (Providone-Iodine) Solution | Penn Veterinary Supply Inc/ Fisher Scien | NC0158124 | |
| Buprenorphine HCL Injection 0.3 mg/mL | Generic/ Covetrus | 59122 | Must obtain state and federal DEA license for purchasing controlled substance |
| Calcein-AM | Sigma Aldrich | 206700-1MG | |
| Cotton Swabs and Applicators | Fisherbrand/ Fisher Scientific | 22-363-172 | |
| Collagenase P | Roche | 11249002001 | Used for digesting bone |
| Cryofilm Type II C(10), 3.5 cm | Section Lab, Japan | CFS 105 | |
| DAPI Staining Solution | Abcam | ab228549 | |
| Digital Gram Scale | Toprime/ Amazon | B06X6LW4V9 | Can use any brand scale to weigh mice |
| Dissecting Forceps | Fisherbrand/ Fisher Scientific | 08-953E; 08-953F | |
| Dissecting Scissors | Fisherbrand/ Fisher Scientific | 08-940 | |
| Endure Low Profile Cryostat/Microtome Blades | Tanner Scientific | TNR315LP | |
| Ethiqa XR Buprenorphine Extended-Release 1.3 mg/mL | Ethiqa XR/ Cevetrus | 72117 | Must obtain state and federal DEA license for purchasing controlled substance |
| Fast Green, 0.2% | Electron Miscroscopy Sciences EMS | 50-319-57 | |
| Fetal Bovine Serum heat inactivated | Gibco/ Thermo Fisher Scientific | A5256801 | Used to make FACS Staining Media (FSM) |
| Fluoromount-G Mounting Solution | SouthernBiotech | 0100-01 | |
| GLUSEAL Topical Skin Adhesive ? | AD Surgical | GLU-550 | |
| Hanks' Balanced Salt Solution (HBSS)¬† | Gibco/ Thermo Fisher Scientific | 14065056 | Used to make FACS Staining Media (FSM) |
| Heating Pad (large) | Boncare/ Amazon | B0CLHSWXSX | Can use any brand heating pad, remove the cloth cover |
| Hematoxylin | MilliporeSigma/ Fisher Scientific | M1051740500 | |
| Hyaluronidase | MilliporeSigma | HX0514 | Used for digesting bone |
| Isoflurane Animal Anesthesia System | VetEquip | 901810 | |
| KUBTEC Parameter 2D Cabinet X-ray System | OncoMed Solutions | N/A | Any X-ray Cabinet imaging system can be used |
| Microcentrifuge Tubes | Corning/ Thomas Scientific | 8600A52 | |
| Microscope Cover Glass, 18mm x 18mm, # 1 Thickness | Globe Scientific | 1414-10 | |
| N-2-hydroxyethylpiperazine-N-2-ethane sulfonic acid HE | Gibco/ Thermo Fisher Scientific | 156300-80 | Used to make FACS Staining Media (FSM) |
| Norland optical Adhesive NO61 | Edmund Optical | 37-322 | Used to adhere cryofilm sections to a slide |
| Propidium Iodide (PI) | Invitrogen/ Thermo Fisher Scientific | P1304MP | Nuclear fluorescent stain used to detect dead cells for vioability counts |
| Rechargeable Electric Trimmer | Wahl/ Amazon | B001FWXKUE | Any clippers can be used, a smaller sized trimmer will make it easier to shave smaller animals |
| Rodent Nose Cone | VetEquip | 921609 | |
| Safranin O, 85% pure, certified | Fisher Scientific | AC419211000 | |
| Seal'N Freeze Cryotray Square Molds | Electron Miscroscopy Sciences EMS | 62642-01 | Used for embedding, other embedding molds can be used. This specific model has a deeper mold, as well as a lid to seal the block |
| Silver Nitrate | Sigma Aldrich | S6506 | silver nitrate for von Kossa stain |
| Sterile Phosphate Buffered Saline (PBS) | Gibco/ Thermo Fisher Scientific | 10010023 | Alternatively, freshly prepared PBS that has been filtered and autoclaved can be used |
| Sterile Standard Scalpels | Integra Miltex/ Fisher Scientific | 12-460-455 | |
| Sucrose Powder 1KG | Thermo Fisher Scientific | 036508-A1 | |
| Superfrost Plus Miscroscope Slides | Fisherbrand/ Fisher Scientific | 1255015 | |
| Sure-Tek Tissue Processing/ Embedding Casettes | Fisherbrand Histosette/ Fisher Scientific | 22-048-141 | |
| Tissue Plus Optimal Cutting Temperature O.C.T Compou | Fisher Scientific | 23-730-571 | |
| Tissue Sponges | Epredia/ Fisher Scientific | 84-53 | Tissue sponges protect tissues from overproocessing |
| Toluidine Blue | Fisher Scientific | AC348600050 | |
